# Low expression of EXOSC2 protects against clinical COVID-19 and impedes SARS-CoV-2 replication

**DOI:** 10.1101/2022.03.06.483172

**Authors:** Tobias Moll, Valerie Odon, Calum Harvey, Mark O Collins, Andrew Peden, John Franklin, Emily Graves, Jack N.G. Marshall, Cleide dos Santos Souza, Sai Zhang, Mimoun Azzouz, David Gordon, Nevan Krogan, Laura Ferraiuolo, Michael P Snyder, Pamela J Shaw, Jan Rehwinkel, Johnathan Cooper-Knock

**Affiliations:** Sheffield Institute for Translational Neuroscience, University of Sheffield, Sheffield, UK; Medical Research Council Human Immunology Unit, Medical Research Council Weatherall Institute of Molecular Medicine, Radcliffe Department of Medicine, University of Oxford, Oxford, UK; School of Biosciences, University of Sheffield, Sheffield, United Kingdom; Department of Genetics, Stanford University School of Medicine, Stanford, CA, USA; Center for Genomics and Personalized Medicine, Stanford University School of Medicine, Stanford, CA, USA; Department of Cellular and Molecular Pharmacology, University of California San Francisco, San Francisco, CA, USA; Quantitative Biosciences Institute (QBI), University of California San Francisco, San Francisco, CA, USA

## Abstract

New therapeutic targets are a valuable resource in the struggle to reduce the morbidity and mortality associated with the COVID-19 pandemic, caused by the SARS-CoV-2 virus. Genome-wide association studies (GWAS) have identified risk loci, but some loci are associated with co-morbidities and are not specific to host-virus interactions. Here, we identify and experimentally validate a link between reduced expression of EXOSC2 and reduced SARS-CoV-2 replication. EXOSC2 was one of 332 host proteins examined, all of which interact directly with SARS-CoV-2 proteins; EXOSC2 interacts with Nsp8 which forms part of the viral RNA polymerase. Lung-specific eQTLs were identified from GTEx (v7) for each of the 332 host proteins. Aggregating COVID-19 GWAS statistics for gene-specific eQTLs revealed an association between increased expression of *EXOSC2* and higher risk of clinical COVID-19 which survived stringent multiple testing correction. EXOSC2 is a component of the RNA exosome and indeed, LC-MS/MS analysis of protein pulldowns demonstrated an interaction between the SARS-CoV-2 RNA polymerase and the majority of human RNA exosome components. CRISPR/Cas9 introduction of nonsense mutations within *EXOSC2* in Calu-3 cells reduced EXOSC2 protein expression, impeded SARS-CoV-2 replication and upregulated oligoadenylate synthase (*OAS)* genes, which have been linked to a successful immune response against SARS-CoV-2. Reduced EXOSC2 expression did not reduce cellular viability. OAS gene expression changes occurred independent of infection and in the absence of significant upregulation of other interferon-stimulated genes (ISGs). Targeted depletion or functional inhibition of EXOSC2 may be a safe and effective strategy to protect at-risk individuals against clinical COVID-19.

## Introduction

Infection with severe acute respiratory syndrome coronavirus 2 (SARS-CoV-2) giving rise to coronavirus disease 2019 (COVID-19) has caused a global pandemic with almost unprecedented morbidity and mortality (Dong et al. 2020). Vaccination efforts have led to early successes (Shilo et al. 2021), but the prospect of evolving variants capable of immune-escape (Darby and Hiscox 2021) and a time-limit on vaccine effectiveness (Pouwels et al. 2021) highlight the importance of efforts to better understand COVID-19 pathogenesis and to develop effective treatments.

SARS-CoV-2 gains entry to host cells in the upper airway via interaction with the cell-surface proteins ACE2 and TMPRSS2 (Hoffmann et al. 2020) but, like all viruses, the process of viral replication requires interaction with a range of host proteins intracellularly. Early work to determine important interactions between viral and host proteins outlined a set of 332 high confidence interactions (Gordon et al. 2020). We hypothesised that variation in function and expression of host proteins involved in these interactions could modify SARS-CoV-2 replication and potentially increase or decrease the risk of symptomatic infection.

To date therapeutic approaches to COVID-19 have focused on modulation of the host immune response. Severe infection is thought to result from a combination of uncontrolled viral replication and a late hyperinflammatory response leading to acute respiratory distress syndrome (ARDS) (Brodin 2021). As a result current therapeutic approaches seek to either boost host immunity (e.g. vaccines) (Zhang et al. 2021; Olliaro et al. 2021), reduce viral replication (e.g. molpunavir (Jayk Bernal et al. 2021)) or reduce hyperinflammation (e.g. dexamethasone (The RECOVERY Collaborative Group. 2021)). Very few strategies have sought to modify a host protein which interacts with the virus: an exception is carmostat, a TMPRSS2 inhibitor, for which there is no good evidence of effectiveness (Gunst et al. 2021).

We performed an unbiased genetic screen using eQTL data from GTEx (Lonsdale et al. 2013) describing gene expression changes in lung tissue. We tested all 332 genes encoding proteins which interact with SARS-CoV-2 proteins, to determine whether gene expression was linked to the risk of clinically symptomatic COVID-19. Increased *EXOSC2* expression within the lung was significantly associated with a higher risk of COVID-19 after stringent multiple testing correction. *EXOSC2* encodes a component of the RNA exosome. We engineered Calu-3 cells to reduce expression of *EXOSC2* and demonstrated a significant suppression of SARS-CoV-2 replication. Transcriptome analysis revealed that reduced expression of EXOSC2 leads to an upregulation of *OAS* gene expression which is independent of infection or inflammation, possibly as part of a homeostatic response (Mullani et al. 2021). OAS proteins are key mediators of viral RNA degradation (Choi et al. 2015) and have been linked to a successful immune response against SARS-CoV-2 (Huffman et al. 2022; Wickenhagen et al. 2021); it is likely that OAS protein upregulation is one reason for the negative effect of reduced EXOSC2 on SARS-CoV-2 replication. Our data suggest that EXOSC2 is a novel therapeutic target for preventing uncontrolled replication of SARS-CoV-2 following infection. Our approach is summarised in **Fig. 1A**.

**Figure 1:**
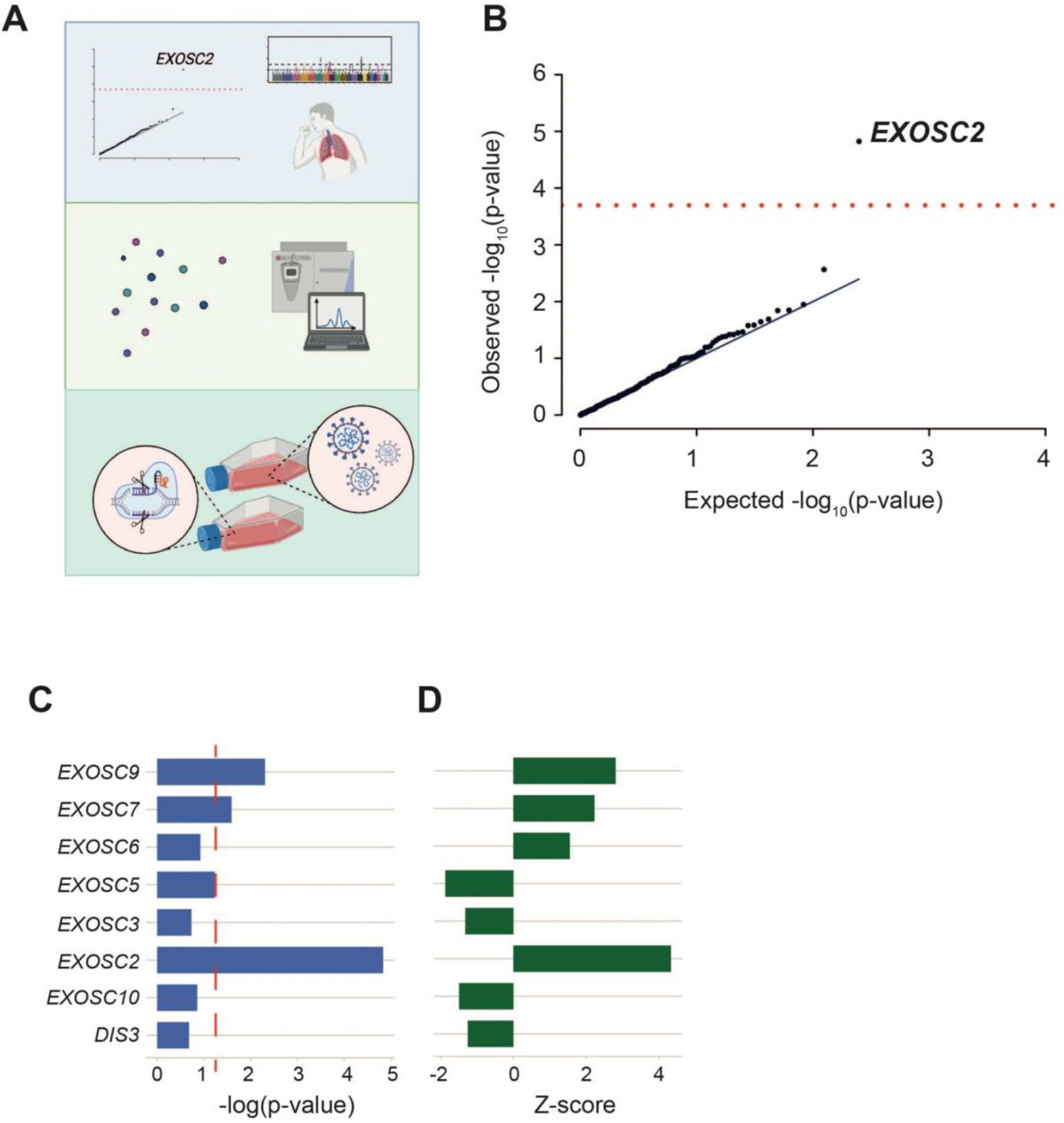
Unbiased screen of host proteins identified as high confidence interacting partners of SARS-CoV-2 proteins links RNA exosome components to risk of clinical COVID-19. (**a**) Schematic of the study design. Known host-viral interactions were screened for disease-association by combining lung-specific eQTLs with a GWAS for COVID-19 symptoms. Identification of a positive correlation between *EXOSC2* expression and increased severity of COVID-19 led to further study of interactions between the SARS-CoV-2 polymerase and the entire human RNA exosome by AP-MS. Finally, CRISPR editing of *EXOSC2* within human lung cells and subsequent infection with SARS-CoV-2 facilitated validation of the relationship between EXOSC2 expression and viral replication and interrogation of the underlying biological mechanism. (**b**) Lung eQTLs were used to group genetic variants according to their effect on expression of 332 host genes encoding proteins which interact with viral proteins. Only expression of *EXOSC2* was significantly associated with clinical risk of COVID-19 after Bonferroni multiple testing (red line). (**c-d**) Lung eQTLs were used to group genetic variants according to their effect on expression of all genes encoding components of the RNA exosome. Expression levels of *EXOSC7, EXOSC9* and *EXOSC2* were significantly linked to clinical COVID-19 and in each case higher expression was associated with higher risk of infection. p=0.05 is indicated by a red dashed line.

## Results

### Unbiased genetic screen highlights RNA exosome components in the defence against SARS-CoV-2

We hypothesised that changes in expression of host proteins which interact with viral proteins could modify the risk of clinical COVID-19. To test this, we focused on genetic variation associated with gene expression changes within lung tissue available from GTEx (v7) (Lonsdale et al. 2013). We measured risk of clinical COVID-19 using a specific set of symptoms (Menni et al. 2020) rather than a positive test to maximise detection of clinically significant COVID-19. Risk of clinical COVID-19 was assigned to specific genetic variants by GWAS (COVID-19 Host Genetics Initiative 2021). We did not focus on hospitalised or severe COVID-19 to minimise confounding by host co-morbidities and immune function, which have been closely associated with COVID-19 mortality (Fathi et al. 2021; Brodin 2021) but may not reflect intracellular interactions between host and viral proteins.

Lung eQTLs were available for 208 of 332 high confidence COVID-19 interacting partners (Gordon et al. 2020). Using this information, we aggregated genetic variants according to their effect on expression of COVID-19 interacting partners. We then used GWAS data to test whether expression changes are associated with higher or lower risk of clinical COVID-19 (**Methods**). After Bonferroni multiple testing correction only *EXOSC2* expression was significantly associated with COVID-19 risk (**Supplementary Table S1, Figure 1b**); higher expression of *EXOSC2* was associated with higher risk of clinical COVID-19 (Z=+4.32, p=1.5E-05). There was no evidence of statistical inflation (λGC1000=0.99).

### Affinity purification confirms interaction of the SARS-CoV-2 viral polymerase with the host RNA exosome

EXOSC2 is bound by Nsp8 (Gordon et al. 2020) which forms part of the SARS-CoV-2 polymerase. We hypothesised that the SARS-CoV-2 RNA polymerase may interact with the entire host RNA exosome complex. The original study of high-confidence interactions between viral proteins and host proteins necessarily included stringent thresholds and may have missed certain interactions. Moreover, the viral polymerase includes Nsp7 in addition to Nsp8 (Hillen et al. 2020). To explore this hypothesis, we performed pulldown experiments using Strep-tagged Nsp8 co-expressed with untagged Nsp7. LC-MS/MS analysis of replicates Strep-Nsp8 and control pulldowns were performed. Statistical analysis of label-free quantification data identified a significant enrichment of EXOSC1, EXOSC2, EXOSC3, EXOSC4, EXOSC5, EXOSC6, EXOSC7, EXOSC8, EXOSC9 and EXOSC10 in Strep-Nsp8 pulldowns (FDR<0.05, permutation test, **Fig. 2A-B, Methods**).

**Figure 2:**
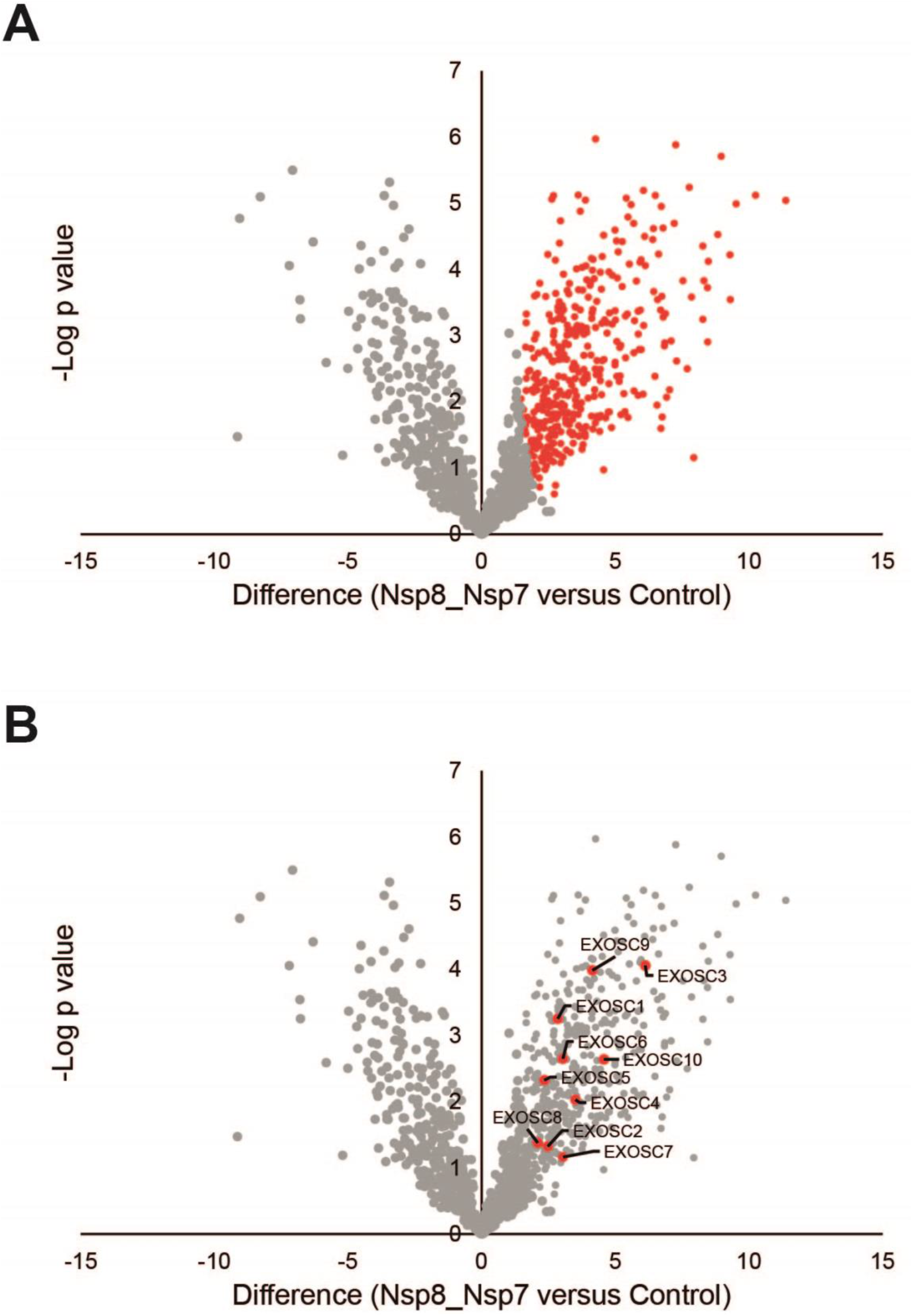
AP-MS analysis confirms the interaction of the SARS-CoV-2 RNA polymerase with EXOSC2 and the majority of components of the host RNA exosome. Replicate affinity purifications of HEK293T cells expressing Strep-Nsp8 and untagged Nsp7 and control purifications (mock-transfected) were analysed by label-free quantitative mass spectrometry. (a) Volcano plot of Strep-Nsp8 pulldowns from cells co-expressing Nsp7 compared to mock-transfected cells. Data points in red are proteins significantly enriched in Strep-Nsp8 pulldowns with a permutation-based FDR 0.05. (b) RNA exosome complex proteins within the set of enriched proteins are labelled.

### Expression of additional RNA exosome components are linked to the defence against SARS-CoV-2

Our immunoprecipitation data strongly suggest that the SARS-CoV-2 polymerase interacts with the entire host RNA exosome complex. In view of this we analysed association between expression of other RNA exosome components with risk for clinical COVID-19. Lung eQTLs were available for *EXOSC2, EXOSC3, EXOSC5, EXOSC6, EXOSC7, EXOSC9, EXOSC10, and DIS3*. Like *EXOSC2*, higher expression levels of *EXOSC7* and *EXOSC9* are significantly associated with higher risk for clinical COVID-19 (p<0.05, **Fig. 1C-D**).

### Infectivity assays reveal reduced viral replication with reduced EXOSC2 expression

We discovered that higher expression of RNA exosome components in lung tissue is associated with higher risk of clinical COVID-19. Next, we sought to provide experimental support for our observations and explore an underlying mechanism. We used CRISPR/Cas9 to introduce loss-of-function mutations within *EXOSC2* in Calu-3 cells. Calu-3 cells are a lung cancer cell line capable of supporting robust SARS-CoV-2 entry and replication (Chu et al. 2020), and a recommended model for viral infection of nasal and bronchotracheal epithelium (Cagno 2020). Calu-3 cells grow in tight monolayers, present villi and are capable of secreting mucins.

We designed a sgRNA to target exon 1 of *EXOSC2*, so as to introduce a series of indels by CRISPR/SpCas9 editing (**Methods**). The efficient introduction of nonsense mutations within edited cells was confirmed by Sanger sequencing and waveform decomposition analysis (**Supplementary Fig. S1A-B**) (Conant et al. 2022). We estimate that ∼60% of alleles were successfully edited (**Supplementary Fig. S1B**); a polyclonal population serves to increase confidence in our downstream analysis. We successfully achieved efficient reduction of *EXOSC2* mRNA levels (63% reduction in mRNA expression, **Supplementary Fig. S1C, Methods**) and protein levels (**Supplementary Fig. S1D, Methods**). The RNA exosome contributes to several RNA processes in the cells, for example it is critical for the production of mature rRNA (Allmang et al. 2000). Reduced expression of EXOSC2 in this cell type was not associated with detectable cell death (MTT assay, **Fig. 3A, Methods**) suggesting that a reduction in RNA exosome expression may be well tolerated in human lung cells. We utilised unedited wild-type (WT) Calu-3 cells, and a commercially available control sgRNA targeting *HPRT* as negative controls.

**Figure 3:**
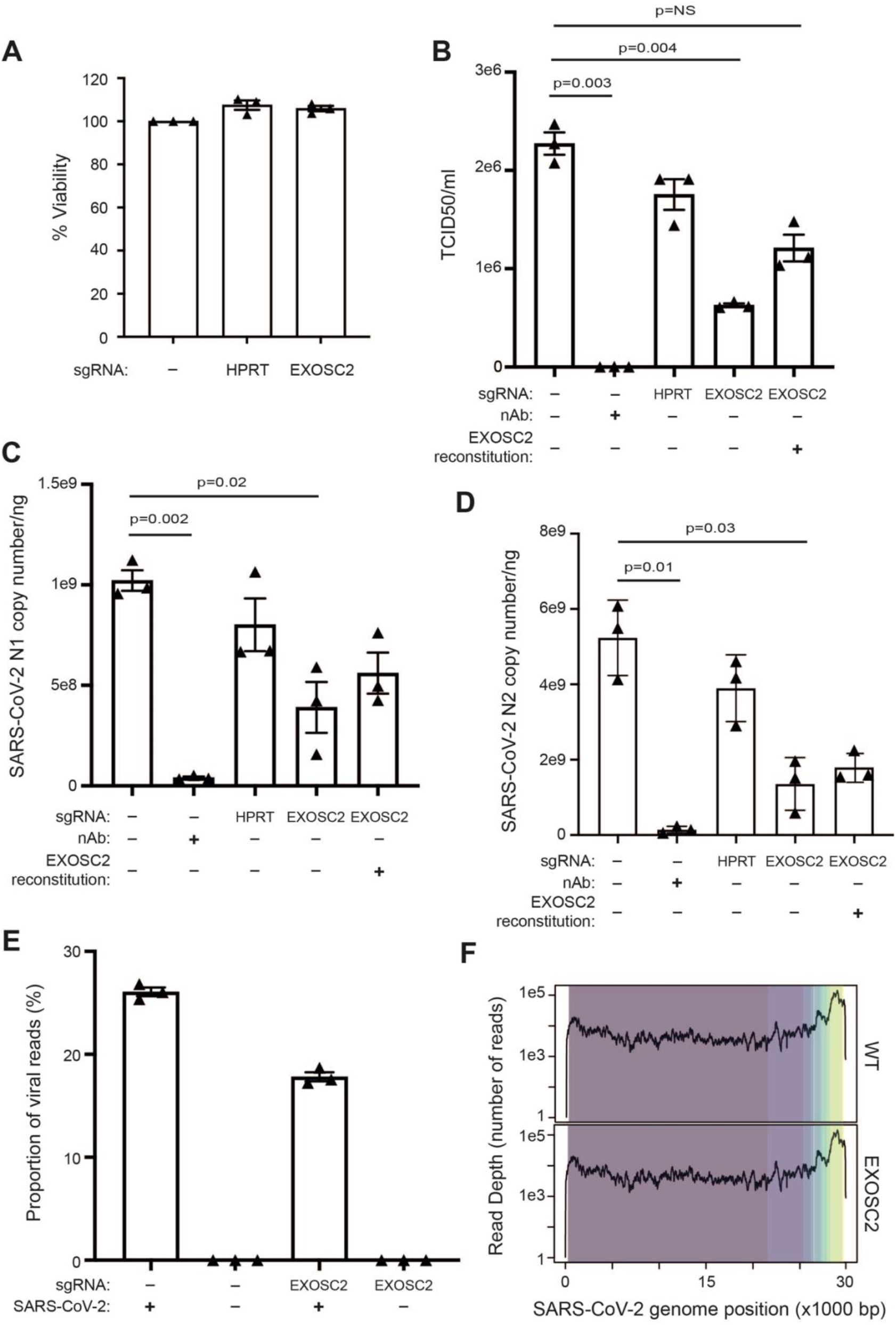
Reduced expression of EXOSC2 in Calu-3 cells is not toxic and leads to reduced viral replication. (**a**) Calu-3 cells were targeted with the indicated sgRNAs and cell viability was analysed by MTT assay. Data for unedited control cells were set to 100%. (**b-f**) Calu-3 cells targeted with sgRNAs and subsequently reconstituted with EXOSC2 as indicated were infected with SARS-CoV-2 (MOI=1) for 17 hours. As a negative control, cells infected with virus were exposed to a neutralising antibody. (**b**) Viral titres in supernatant samples were analysed by TCID50 assay. (**c-d**). Viral RNA levels were measured by absolute RT-qPCR quantification of N1 and N2 SARS-CoV-2 genomic RNA. (**e**) Viral genomic reads as a proportion of total RNA-sequencing reads. (**f**) Viral genomic RNA-sequencing reads mapped across the SARS-CoV-2 genome by normalised read-depth; colours represent distinct viral transcripts. Data are from three independent biological repeats. In panels (**a**-**e**), individual data points are shown with mean and standard error. Significance was tested by paired t-test and p values are indicated.

CRISPR edited Calu-3 cells and control cells were infected with SARS CoV-2 strain Victoria using a multiplicity of infection (MOI) of 1. We confirmed viral replication by absolute RT-qPCR quantification of viral genomes and by the median tissue culture infectious dose (TCID50) (**Methods**). As a positive control we utilised a neutralising antibody for SARS-COV-2 (**Methods**). To achieve absolute quantification of viral genomes we used two nucleocapsid gene targets, N1 and N2 (Holshue et al. 2020) (**Methods**). As predicted, reduced expression of EXOSC2 was associated with a significant reduction in viral infectivity (72% reduction, p=0.004, paired t-test, **Fig. 3B**) and in viral genome replication (N1: 62% reduction, p=0.02; N2: 74% reduction, p=0.03; paired t-test, **Fig. 3C-D**) compared to infection of WT unedited Calu-3 cells.

As a positive control we performed reconstitution of EXOSC2 by overexpression of a sgRNA-resistant plasmid encoding EXOSC2 (**Methods**). Reconstitution was confirmed by immunoblotting (**Supplementary Fig. 1d**). Reconstitution of EXOSC2 followed by infection with SARS-CoV-2 led to increased viral infectivity (93% increase, **Fig. 3B**) and replication (N1: 44% increase; N2: 32% increase; **Fig. 3C-D**) although these changes were not statistically significant. The failure of complete recovery of viral infectivity and replication may be because both EXOSC2 depletion and reconstitution were variable across the population of cells and/or because reduced expression of EXOSC2 changes cellular homeostasis in such a way that is not easily reversible.

### Transcriptome analysis links reduced EXOSC2 function to reduced SARS-CoV-2 replication and increased expression of OAS proteins

Infectivity assays confirmed our hypothesis based on genetic analyses, that reduced EXOSC2 function is protective against SARS-CoV-2 infection. To gain further insight into the biological mechanism underpinning our observations we analysed changes in the total transcriptome of Calu-3 cells in the presence/absence of SARS-CoV-2 infection, both with and without CRISPR editing of *EXOSC2*. We performed RNA-sequencing using three biological repeats for each condition (**Methods**).

Mapping of RNA-sequencing reads to the SARS-CoV-2 genome (**Methods**) confirmed our previous finding that there are significantly fewer viral reads in the presence of reduced EXOSC2 expression (**Fig. 3E**). To determine whether this was due to differential expression of viral subgenomic RNAs we examined read-depth across the SARS-CoV-2 genome; there was no significant difference in the presence/absence of reduced EXOSC2 expression (**Fig. 3F**). We compared the ratio of reads mapped to Orf1 compared to Orf10 as a measure of transcription of subgenomic RNA (**Methods**) but there was no significant difference between the sample groups (t-test, p=0.14). Overall, we conclude that reduced expression of EXOSC2 impacts overall viral replication rather than altering expression of specific viral transcripts.

Next we analysed RNA-sequencing reads mapped to the human genome (**Methods**). Principal components analysis revealed good consistency between biological replicates (**Fig. 4A**). As expected, the first principal component and the largest change in gene expression, was associated with the presence or absence of SARS-CoV-2 infection. The most significant functional enrichment by adjusted p-value, within the set of 2,445 genes up-regulated in WT Calu-3 cells after SARS-CoV-2 infection (**Supplementary Fig. S2A**), was the KEGG pathway ‘TNF signalling pathway’ (Fisher exact test, adjusted p=2.28e-21, OR=7.87) suggesting that this represents the immune response to acute SARS-CoV-2 infection. Our observations are consistent with previous literature: the 2,445 genes were highly enriched with reported gene expression changes in Calu-3 cells infected with SARS-CoV-2 (Fisher exact test, adjusted p=9.44e-245, OR=27.78) (Wyler et al. 2021)). We examined expression of these 2,445 genes across all conditions and concluded that the overall cellular response to SARS-CoV-2 infection was independent of *EXOSC2* gene editing (**Fig. 4B, Supplementary Fig. S2A-B**). Indeed, *IL6*, a key inflammatory gene, which is upregulated in patients suffering COVID-19 (Manjili et al. 2020), was upregulated in the context of SARS-CoV-2 infection (p-value = 6.44e-73, FC = 3.32) but downregulated in uninfected EXOSC2 edited Calu-3 cells compared to unedited cells (p=3.45e-10, FC = 0.67, **Fig. 4C**). Reduced EXOSC2 expression within infected Calu-3 cells produced 903 differentially expressed genes (**Supplementary Fig. 2C**) which represents the effects of EXOSC2 depletion combined with reduced viral replication (**Fig. 3E**).

**Figure 4:**
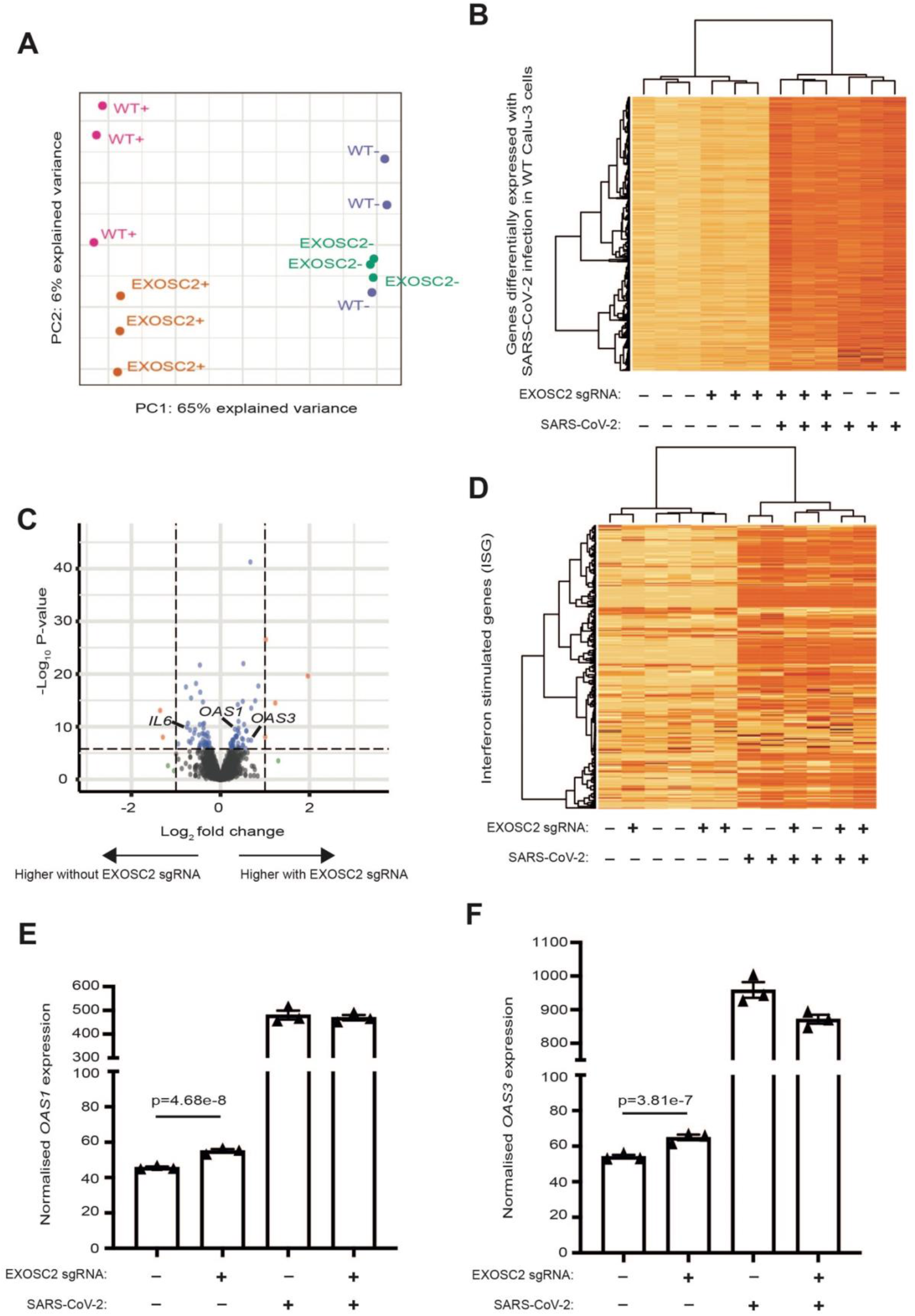
Transcriptomic analysis confirmed the inflammatory response to SARS-CoV-2 infection of Calu-3 cells and identified upregulation of *OAS* genes in the context of reduced EXOSC2 expression. RNA for sequencing was extracted from Calu-3 cells in the presence and absence of CRISPR editing with sgRNA targeted against EXOSC2; with and without infection with SARS-CoV-2 (MOI=1) at 17h; three biological replicates were obtained for all conditions. (**a**) First and second principal components for total gene expression across all sequenced samples. Samples include WT unedited Calu-3 cells and EXOSC2 edited Calu-3 cells; +/- indicates the presence/absence of SARS-CoV-2 infection. (**b**) Heatmap representation of genes upregulated in WT cells in the presence of SARS-CoV-2 infection. A darker colour indicates higher expression. (**c**) Volcano plot to compare gene expression in uninfected Calu-3 cells with and without CRISPR editing of EXOSC2. Dotted lines represent fold change of +/- 2 and a Bonferroni multiple testing threshold for p-value by genewise exact test. (**d**) Heatmap representation of 397 ISGs (Schoggins et al. 2011) across all sequenced samples. (**e-f**) Normalised expression of OAS1 (**d**) and OAS3 (**e**) in all four conditions.

Next we examined changes in gene expression in Calu-3 cells with reduced EXOSC2 expression prior to SARS-CoV-2 infection. In uninfected cells there were 364 differentially expressed genes as a result of EXOSC2 depletion (**Fig. 4C**). This set of genes was not enriched with ‘TNF signalling pathway’ genes (p=0.68), unlike the response to SARS-CoV-2 infection. Unexpectedly, the most significant functional enrichment by adjusted p-value, within this gene set was the KEGG pathway ‘Coronavirus disease’ (Fisher exact test, adjusted p-value=7.7e-4, OR=4.64) which included upregulation of *OAS1* (genewise exact test (Robinson and Smyth 2008), p=4.68e-8, FC=1.21, **Fig. 4E**) and *OAS3* (p=3.81e-7, FC=1.20, **Fig. 4F**). Both *OAS1* and *OAS3* genes encode enzymes which activate ribonuclease L to degrade intracellular double-stranded RNA as part of the antiviral response (Choi et al. 2015) and cellular homeostasis (Mullani et al. 2021).

*OAS* genes form part of the set of interferon-stimulated genes (ISG) (Schoggins et al. 2011). We wondered whether EXOSC2 knockdown produced a non-specific upregulation of ISGs, perhaps via increased concentration of dsRNAs. Of 397 ISGs, only 14 genes were differentially expressed in uninfected Calu-3 cells after CRISPR editing of EXOSC2 which is less than would be expected by chance at a 5% significance level. A heatmap of ISGs across all four conditions (**Fig. 4D**) demonstrates that ISGs are upregulated following SARS-CoV-2 infection but do not efficiently separate cell lines based on EXOSC2 editing status. Overall we conclude that the observed upregulation of OAS genes in the context of reduced EXOSC2 expression is specific and not associated with activation of all ISGs.

## Discussion

The global COVID-19 pandemic entered a new stage when the long-term effectiveness of vaccines was first questioned (Pouwels et al. 2021). New therapeutic strategies are required to prevent COVID-19 infection and associated morbidity and mortality. Our study is data-driven; we have harnessed the power of large-scale GWAS based on real-world observation to identify factors which reduce the risk for clinical COVID-19 in human patients. However, we have also taken advantage of insights at a molecular level to identify candidate host-virus interactions which may influence SARS-CoV-2 replication (Gordon et al. 2020). By validating our work using live virus we have demonstrated the validity of our findings to reduce viral replication.

The RNA exosome functions in RNA quality control; aberrant or unwanted RNAs are captured and degraded by the complex (Kilchert et al. 2016). EXOSC2 and EXOSC3 form part of a cap structure with RNA-binding activity responsible for passing substrate RNAs into a barrel-like structure formed by components including EXOSC5 and EXOSC8, from which RNAs then access the catalytically active core. RNA degradation is performed by EXOSC10, DIS3 or DIS3L; the precise catalytic subunit varies by subcellular location (Tomecki et al. 2010). The RNA exosome has been implicated in the antiviral response (Molleston and Cherry 2017) and has been observed to associate with other RNases.

A key question is the mechanism underlying the protection we have observed in cells and patients with reduced EXOSC2 expression. While we have not conclusively answered this question we highlight a number of possibilities. Previously, SKIV2L, a component of the RNA exosome-activating SKI complex, was shown to limit baseline type I IFN responses, which are induced by RNA sensors in settings of *SKIV2L* deficiency (Eckard et al. 2014). Moreover, pharmacological inhibition of the SKI complex limits SARS-CoV-2 replication (Weston et al. 2020). We did not observe broad ISG induction in EXOSC2-depleted cells, suggesting that reduced SARS-CoV-2 replication was not simply a consequence of elevated baseline ISG expression in these cells.

We identified EXOSC2 because it interacts with the SARS-CoV-2 polymerase (Gordon et al. 2020). Interaction between the host RNA exosome and the viral RNA polymerase is important for viral replication for influenza A virus (IAV) and Lassa virus (Ho et al. 2021). These viruses do not encode their own capping enzymes and so take advantage of host caps which are a byproduct of RNA degradation by the RNA exosome. Loss of RNA exosome function has been previously shown to protect against IAV infection (Rialdi et al. 2017). However, current knowledge regarding coronaviruses and SARS-CoV-2 in particular, indicates that this virus encodes its own mRNA capping machinery (Viswanathan et al. 2020), suggesting that this is not the mechanism underpinning our observations. Moreover, we did not observe a difference in production of specific viral transcripts in the presence of reduced EXOSC2 expression, contrary to what might be expected with failure of viral mRNA capping.

Finally, the RNA exosome functions in degradation of exogenous and endogenous RNAs alongside OAS proteins; indeed reciprocal upregulation of OAS proteins has been observed in the context of EXOSC3 depletion (Mullani et al. 2021). The physical association between SARS-CoV-2 and the RNA exosome suggest that the virus is relatively protected against degradation by the exosome, but its vulnerability to OAS proteins is well described (Huffman et al. 2022; Wickenhagen et al. 2021). We observed a modest upregulation of *OAS* transcripts in the context of reduced EXOSC2 expression in the absence of SARS-CoV-2 infection. Consistent with our observations, upregulation of OAS proteins would be expected to reduce viral replication in cells with reduced EXOSC2 expression. It is therefore possible that changes in OAS protein expression contribute to the mechanism linking EXOSC2 expression and SARS-CoV-2 replication, although we have not tested here whether manipulation of OAS proteins modulates the effect of EXOSC2 on viral replication.

Upregulation of the p46 splice variant of *OAS1* has been associated with protection against severe COVID-19 (Huffman et al. 2022). Production of the p46 variant is contingent upon a G allele for SNP rs10774671 whereas Calu-3 cells are homozygous A (Iida et al. 2021) and therefore cannot express the p46 variant. It follows that, if the reduced viral replication we observed in Calu-3 cells expressing low levels of EXOSC2 is mediated via OAS protein upregulation, then it must be achieved only through OAS3 and not OAS1. We conclude that our study of Calu-3 cells may therefore underestimate the effect of OAS protein upregulation on viral replication, in comparison with our genetic study which focused on a population where OAS1 p46 expression will be frequent.

Our genetic study suggests that reduced expression of EXOSC2 is well tolerated in a significant proportion of the population who are relatively immune to clinical COVID-19. Indeed, we note that our genetically edited cells did not show excess toxicity. Future structural biology work will be necessary to determine the mechanism linking the interaction between SARS-CoV-2 and the host RNA exosome and changes in viral replication, however we have suggested candidate mechanisms based on our analyses. We have identified a new therapeutic target with the potential to protect against COVID-19. We anticipate that our work will lead to new understanding and new therapies.

## Acknowledgements

This work was supported by the National Institutes of Health (CEGS 5P50HG00773504, 1P50HL083800, 1R01HL101388, 1R01-HL122939, S10OD025212, P30DK116074, and UM1HG009442 to M.P.S.), the Wellcome Trust (216596/Z/19/Z to J.C.-K.), the NIHR (NF-SI-0617-10077 to P.J.S.), the BBSRC (BB/S009566/1 to A.A.P and M.O.C), and the UK Medical Research Council (MRC core funding of the MRC Human Immunology Unit to J.R.). C.H. is supported by a studentship from the MND Association (899-792). We also acknowledge support from a Kingsland fellowship (T.M.), and the NIHR Sheffield Biomedical Research Centre for Translational Neuroscience (IS-BRC-1215-20017). We thank the Oxford Genomics Centre at the Wellcome Centre for Human Genetics (funded by Wellcome Trust grant reference 203141/Z/16/Z) for the generation and initial processing of sequencing data. Figure 1A was created with BioRender.com. Neutralising antibody was a kind gift from Alain Townsend.

## Author Contributions

TM, JR, VO and JCK conceived and designed the study. TM, VO, CH, MOC, AP, JF, EG, JNGM, CDSS, SZ, MA, DG, NK, LF, MPS, PJS, JR and JCK were responsible for data acquisition. TM, VO, CH, MOC, JF, EG, JNGM, CDSS, SZ, MA, JR and JCK were responsible for analysis of data. TM, VO, CH, MOC, JF, EG, JNGM, CDSS, SZ, MA, JR and JCK were responsible for interpretation of data. TM, VO, CH, MOC, SZ, JR and JCK prepared the manuscript with assistance from all authors. All authors meet the four ICMJE authorship criteria, and were responsible for revising the manuscript, approving the final version for publication, and for the accuracy and integrity of the work.

## Methods

### Association of lung gene expression with risk of COVID-19

Association testing was performed using FUSION (Gusev et al. 2016) using pre-computed weights for lung gene expression from GTEx (v7) (Lonsdale et al. 2013). All eQTLs were used regardless of significance. We used the ANA7 phenotype from DF2 which included 1,294 individuals with clinical evidence of COVID-19 and 26,969 asymptomatic controls.

### Affinity purification of Nsp8-associated complexes and mass spectrometry analysis

HEK293T cells were transfected with Strep-Nsp8 (Gordon et al. 2020) and untagged Nsp7 using PEI for 24 hours. Mock transfected cells were used for control purifications. Cell pellets were collected, proteins lysates were prepared, and affinity purification was performed using 4 biological replicates of each group as previously described (Gordon et al. 2020). Elution of affinity-purified proteins was performed by incubation of MagStrep “type3” beads with 50 μl of elution buffer (5% SDS, 50 mM Tris, pH 7.4) at 70°C for 15 minutes. Protein reduction was performed by adding TCEP to a final concentration of 5 mM and incubation at 70°C for 15 minutes and alkylation by adding iodoacetamide to a final concentration of 10 mM and incubation at 37°C for 30 minutes. Sample clean-up was performed using Suspension trapping (S-Trap) according to the manufacturer’s instructions (PROTIFI). Tryptic digestion was performed by adding 1 μg of trypsin (Pierce, sequencing grade) and incubating at 47°C for 60 minutes. Eluted peptides were dried to completion in a vacuum concentrator (Eppendorf).

Samples were re-suspended in 40 μl of 0.5% formic acid, and 18 μl was analysed by nanoflow LC-MS/MS using an Orbitrap Elite (Thermo Fisher) hybrid mass spectrometer equipped with a nanospray source, coupled to an Ultimate RSLCnano LC System (Dionex) at the biOMICS facility at the University of Sheffield. The system was controlled by Xcalibur 3.0.63 (Thermo Fisher) and DCMSLink (Dionex). Peptides were desalted on-line using an Acclaim PepMap 100 C18 nano/capillary BioLC, 100A nanoViper 20 mm x 75 µm I.D. particle size 3 µm (Fisher Scientific) at a flow rate of 5 μl/min and then separated using a 125-min gradient from 5 to 35% buffer B (0.5% formic acid in 80% acetonitrile) on an EASY-Spray column, 50 cm × 50 μm ID, PepMap C18, 2 μm particles, 100 Å pore size (Fisher Scientific) at a flow rate of 0.25 μl/min. The Orbitrap Elite was operated with a cycle of one MS (in the Orbitrap) acquired at a resolution of 60,000 at m/z 400, with the top 20 most abundant multiply charged (2+ and higher) ions in a given chromatographic window subjected to MS/MS fragmentation in the linear ion trap. An FTMS target value of 1e^6^ and an ion trap MSn target value of 1e^4^ were used with the lock mass (445.120025) enabled. Maximum FTMS scan accumulation time of 100 ms and maximum ion trap MSn scan accumulation time of 50 ms were used. Dynamic exclusion was enabled with a repeat duration of 45 s with an exclusion list of 500 and an exclusion duration of 30 s.

### Mass-spectrometry data analysis

Raw data files were processed using MaxQuant (Version 1.6.10.43) (Tyanova et al. 2016). Data were searched against a combined human and SARS-CoV-2 UniProt sequence database (Dec 2019) using the following search parameters: digestion set to Trypsin/P with a maximum of 2 missed cleavages, oxidation (M), N-terminal protein acetylation as variable modifications, cysteine carbamido-methylation as a fixed modification, match between runs enabled with a match time window of 0.7 min and a 20-min alignment time window, label-free quantification was enabled with a minimum ratio count of 2, minimum number of neighbours of 3 and an average number of neighbours of 6. A first search precursor tolerance of 20ppm and a main search precursor tolerance of 4.5 ppm was used for FTMS scans and a 0.5 Da tolerance for ITMS scans. A protein FDR of 0.01 and a peptide FDR of 0.01 were used for identification level cut-offs.

Protein group output files generated by MaxQuant were loaded into Perseus version 1.6.10.50. The matrix was filtered to remove all proteins that were potential contaminants, only identified by site and reverse sequences. The LFQ intensities were then transformed by log2(x), normalised by subtraction of the median value, and individual intensity columns were grouped by experiment. Proteins were filtered to keep only those with a minimum of 3 valid values in at least one group. The distribution of intensities was checked to ensure standard distribution for each replicate. Missing values were randomly imputed with a width of 0.3 and downshift of 1.8 from the standard deviation. To identify significant differences between groups, two-sided Student’s t-tests were performed with a permutation-based FDR of 0.05. The mass spectrometry data have been deposited to the ProteomeXchange Consortium via the PRIDE partner repository with the dataset identifier PXD031611.

### Calu-3 cell culture

Calu-3 cells were cultured in Dulbecco’s Modified Eagle’s Medium (DMEM)/F12 (1:1) GlutaMAX™ (ThermoFisher Scientific) supplemented with 10% FBS, 1% NEAA, 1% sodium pyruvate, 1% penicillin-streptomycin and maintained at 37°C, 5% CO_2_ and passaged with TrypLE™ Express 1X (ThermoFisher Scientific) when ∼80% confluent. All experimental work was performed on cells within the range of 20-30 passages.

### CRISPR-Cas9 editing

A sgRNA targeting exon 1 of EXOSC2 (5’-GAUACAAUCACUACGGACAC-3’) was designed using the CRISPOR tool (http://crispor.tefor.net/) (Concordet & Haeussler, 2018). Design was guided by available protospacer adjacent motifs and predicted on- and off-target efficiencies. A validated, commercially available sgRNA targeting HPRT (IDT) was used as a CRISPR control. sgRNA duplexes were assembled from tracrRNA and crRNA in a thermocycler according to manufacturer’s instructions under RNase-free conditions. Cells were cultured to ensure 70%–90% confluency on the day of transfection. 24-well plates containing 500 µL of antibiotic-free (DMEM)/F12 (1:1) GlutaMAX™ (ThermoFisher Scientific) were incubated at 37°C. CRISPR/Cas9 Ribonucleoproteins were formed by complexing 240ng sgRNA duplex with 1250ng Alt-R V3 Cas9 Protein (IDT) in 10 μL buffer R (Neon™ Transfection System 10 μL Kit, ThermoFisher Scientific) - a 1:1 molar ratio - for 10 minutes at room temperature (RT). 100,000 viable cells were aliquoted per transfection and centrifuged at 400 x g for 5 minutes at RT. Cells were washed in calcium- and magnesium-free Dulbecco’s Phosphate Buffered Saline (Sigma) and centrifuged at 400 x g for 5 minutes at RT. Cell pellets were resuspended in 10 μL buffer R containing Cas9 protein and sgRNA duplexes. 2 μL of 10.8 μM electroporation enhancer (IDT) was added and the solution mixed thoroughly to ensure a suspension of single cells. 10 μL of this mixture was loaded into a Neon transfection system (ThermoFisher Scientific) and electroporated according to manufacturer’s instructions (1400V, 2 pulse, 20 s pulse width). Cells were then transferred to pre-warmed media in 24-well plates. To assess editing efficiency genomic DNA was isolated from CRISPR-edited and control cells using a GenElute Mammalian DNA Miniprep Kit (Sigma) according to manufacturer’s instructions. A ∼400bp region around the expected Cas9 cut site in exon1 of EXOSC2 was amplified by PCR using PfuTurbo DNA polymerase (Agilent), according to the manufacturer’s instructions using primers: AGGCCGTGAGTTCTCATTGG (fwd) and GGGTTTCAGGGAGCTGAGAC (rvs) (Sigma-Aldrich). Expected amplification was confirmed using gel electrophoresis, and the products were Sanger-sequenced (Source BioScience). Sequencing trace files were uploaded to TIDE (http://shinyapps.datacurators.nl/tide/) and ICE (https://ice.synthego.com), and an indel efficiency calculated.

### MTT assay

Cell viability was assessed using the MTT Assay kit (Abcam) according to the manufacturer’s instructions. Viable cells but not dead cells can metabolise MTT (3-(4,5-dimethylthiazol-2-yl)-2,5-diphenyltetrazolium bromide) to an insoluble formazan product; when solubilised, the compound can be read at OD590 nm. The measured absorbance is proportional to the number of viable cells. 60,000 Calu-3 cells were seeded in each well of a 96-well plate. After 48 hours, the culture medium was discarded and 50 µl of fresh serum-free medium and 50 µl MTT reagent was added to each well, including to a background (no cells) control. The plates were returned to the 37°C incubator for 3 hours before the MTT/media solution was discarded and 150 µl of MTT solvent was added to each well. The plates were incubated for a further 15 minutes in foil on a hula mixer and the absorbance recorded at 590 nm. The replicate values were corrected for background and averaged.

### Cloning and viral transduction

The human *EXOSC2* open-reading frame was amplified from HEK293 cDNA using oligonucleotides: gctagcATGGCGATGGAGATGAGGC (fwd) and ctcgagTCCCTCCTGTTCCAAAAGCCT (rvs) and cloned as a NheI/XhoI PCR fragment into the NheI/XhoI restriction sites of a lentiviral self-inactivating transfer vector (SIN) containing a woodchuck hepatitis virus post regulatory element (W) to overexpress EXOSC2 under a PGK promoter (pLV_SIN-W-PGK-EXOSC2). All plasmids were validated by Sanger-sequencing and sequences are available upon request.

HEK293T cells were used for lentiviral production, plated at a density of 3 × 10^6^ per 10 cm dish. Cells were transfected using a calcium chloride transfection containing 0.5M calcium chloride (Sigma), 2X HEPES Buffered Saline (Sigma) and four lentiviral component plasmids; pCMV delta 8.2 (13 µg), pRSV-Rev (3 µg), pMD.G (3.75 µg) (Addgene) and pLV_SIN-W-PGK-EXOSC2 (13 µg) (Delgon et al., 2000). Transfection mix added dropwise to each plate and left overnight, with a full media change carried out the following morning. Cells were incubated for a further 48 hours before all media was collected and filtered using a 0.45 µm filter (Sigma). Equally loaded tubes (Beckman) were then spun at 19,000rpm/90 minutes/4°C using an ultracentrifuge and a SW28 hanging rota (Beckman). All supernatant was removed and each viral pellet was resuspended with 300 µl of 1% Bovine Serum Albumin (Tocris Bioscience) in Phosphate Buffer Solution (Sigma). Each tube was incubated on ice for 1 hour and then combined into one homogeneous solution before being aliquoted and stored at −80°C.

Viral titres were measured through qPCR against a virus of known biological titre (FACS titration). Genomic DNA was isolated from cells (described previously) which had been transduced with a serial dilution of virus. Viral genomic integration was measured using WPRE primers: CCCGTACGGCTTTCGTTTTC (fwd) and CAAACACAGAGCACACCACG (rvs).

### SARS-CoV-2 production

SARS-CoV-2 strain Victoria was produced by infecting Vero E6 cells at an MOI of 0.01 in DMEM supplemented with 2% FCS for 72 hours, until cytopathic effects were visible. The supernatant containing viral particles was harvested, aliquoted and stored at −80°C. The titre of the SARS-CoV-2 stock was determined by plaque assay using Vero E6 cells.

### SARS-CoV-2 challenge

1 million Calu-3 cells were seeded in each well of a 12-well plate. Cells were left to adhere for 24 hours. The cells were infected with SARS-CoV-2 strain Victoria at MOI 1 in 2% FCS-medium, allowing a total volume of 1ml per well. The cells were returned to the incubator and harvested at 17 hours post-infection when 20% of cells showed cytopathogenic effects (CPE). To neutralise SARS-CoV-2, 1ml of SARS-CoV-2 was incubated for 1h before infection with 15µl of antibody clone EY11A with regular mixing. EY11A inhibits SARS-CoV-2 activity by binding strongly to its spike protein. At the end point, the supernatant was collected and preserved at −80°C for future determination of virus titre and following two washes with PBS, RNA was extracted from cells using RNeasy Mini kit (Qiagen) following manufacturer’s instructions.

For the RNA sequencing experiment, 2 million cells were seeded in 6-well plates and after 24 hours, cells were infected with SARS-CoV-2 Victoria strain at MOI 1. At 17 hours post infection cells were washed twice with PBS and harvested for RNA extraction using Qiagen RNEasy mini kit. The quality of the RNA was assessed by nanodrop ratios and Tapestation 4200 (Agilent). RNAs with 260/230 ratio>2 and RIN>9.8 were submitted for library preparation and sequencing.

### TCID50 titration of virus after challenge

96-well plates were seeded with 15,000 Vero E6 cells per well 24 hours before adding virus in 120 µl of 2% FCS medium per well. 40 µl/well of preserved supernatant was added to the monolayer of Vero E6 cells in column 1 of the plate(rows A to H); this provides 8 replicates to determine TCID50/ml. The virus was mixed by carefully pipetting 6 times before changing tips at each column and transferring 40 µl to the next column, performing a serial dilution (1:4) across the plate. The plates were incubated at 37°C for 72 hours and stained with crystal violet. TCID50/ml was determined visually by recording cell death at each dilution and deriving the titre using the freely available online software (https://www.klinikum.uni-heidelberg.de/fileadmin/inst_hygiene/molekulare_virologie/Downloads/TCID50_calculator_v2_17-01-20_MB.xlsx).

### Quantification of SARS-CoV-2 RNA copy number

SARS-CoV-2 RNA copy number was quantified in RNA samples extracted from infected cells using the QuantiTect probe RT-PCR Kit (Qiagen). TWe prepared a standard curve using the ORF9 nucleoprotein coding sequence cloned in pcDNA3. 10µg of plasmid was linearised with Apa1 at 37°C for 2 hours. The DNA was purified with the PCR clean up kit from Qiagen and 1µg of DNA was used in a T7 transcription reaction using Promega T7 RiboMAX (P1320) at 37°C for 30 minutes. The transcribed RNA was cleaned using Zymo RNA clean and concentrator 25 (ZymoResearch-R1017). The copy number was derived using an online calculator (www.nebiocalculator.neb.com ssRNA mass to moles converter).

The RNA standard curve was prepared from diluting RNA template from 10e8 to 10e1 in RNase-free water. RNA samples were diluted 1:5000 and 1µl of standard or sample was dispensed into each well of a 384 well plate to perform the RT-PCR step following manufacturer’s instructions. Quantitation of copy number was performed using N1 and N2 standard probes for SARS-CoV-2 (IDTDNA).

### EXOSC2 reconstitution

5 µg of pLenti-EXOSC2, with 2.5 µg pR8.91 and 2.5 µg of VSV-G plasmids were transfected in a 15 cm dish of Hek293T cells at 80% confluency. The transfection mix was prepared using Fugene 6 (Promega) at 3µl/ug DNA in 500µl (total volume) of Opti-MEM (Thermo Fisher). The transfection mix was incubated following manufacturer’s recommendations and added dropwise to the dish. After 24 hours the medium was changed to 15ml 2%FCS medium. Following a further 24 hours incubation, the supernatant was collected, spun at 1500 rpm to remove cellular debris and passed through 0.45 µM syringe filter (Sartorius). The supernatants were preserved at −80°C.

To transduce cells with EXOSC2-lentivirus, a T25 of EXOSC2 CRISPR-edited cells was incubated for 48 hours with 2 ml of culture media supplement with 1 ml of viral supernatant and 10µg of polybrene. The cells were then returned to normal propagation media.

### Immunoblotting for EXOSC2

500,000 Calu-3 cells were seeded in 12-well plates and left to adhere for 24 hours, the media was discarded, cells were subsequently washed with PBS and 250 µl of RIPA buffer (10 mM TRIS–HCl pH 8, 140 mM NaCl, 1% Triton-X100, 0.1% SDS, 0.1% sodium deoxycholate, 1 mM EDTA, 1 mM EGTA) containing protease inhibitor (Roche-04693132001) was added to the cells. Cells were scraped from wells, dispensed into an Eppendorf tube and agitated for 20 minutes at 4°C, then spun at 13000 rpm for 20 minutes at 4°C. The supernatant was collected and 100µg of total protein (determined by qubit) was supplemented with Laemmli buffer, boiled for 5 minutes and resolved on 4 to 20% TGX mini protean gels (Bio-Rad). The proteins were transferred to PVDF membranes using BioRad Trans-Blot semi-dry transfer. The membrane was blocked for 1 hour with constant rotation in 2.5% Milk-PBS 0.1% tween20 and incubated overnight at 4°C with the EXOSC2 antibody (Proteintech-66099-1-Ig) or GAPDH antibody (Cell Signaling-14C10) diluted in 2.5% Milk-PBS-0.1% Tween20. After 3 washes of 15 minutes each in PBS-0.1%Tween20, the membranes were incubated with secondary antibodies for 1 hour and following 3 further washes with PBST, the membrane was transferred to a fresh falcon tube and incubated with 1ml of Western LightningPlus-ECL chemiluminescent reagent (PerkinElmer). Proteins bands were visualised with an iBright instrument (ThermoFisher). The intensity of EXOSC2 band against GAPDH loading control was assessed using ImageJ.

### qRT-PCR for EXOSC2

Calu-3 cells were lysed on ice using Tri Reagent (Sigma) for 5 minutes in RNase-free conditions. Total RNA was extracted using a Direct-zol RNA Miniprep Kit (Zymo) according to the manufacturer’s instructions. RNA concentration was determined using a NanoDrop spectrophotometer (ThermoFisher Scientific). 2 µg total RNA was converted to cDNA by adding 1 µL 10 mM dNTPs, 1µL 40 µM random hexamer primer, and DNAse/RNAse-free water to a total reaction volume of 14 μL. The mixture was heated to 70°C followed by a 5 minute incubation on ice. 4 μL 5x first strand buffer, 2 μL 0.1 M DTT, and 1 μL M-MLV reverse transcriptase (ThermoFisher Scientific) were then added and cDNA conversion was performed in a thermocycler (37°C for 50 minutes, 70°C for 10 minutes). cDNA was amplified using RT-PCR with Brilliant III SYBR Green (Agilent) as per the manufacturer’s instructions using primers: AACCTGGAGCCTGTCTCTCTT (fwd) and TGATCTGATGTGGAAGGGATGC (rvs). CT analysis was performed using CFX Maestro software (BioRad).

### RNA-sequencing

RNA was extracted from Calu-3 cells in the presence and absence of *EXOSC2* gene editing at 17 hours post SARS-CoV-2 infection at MOI 1. Extracted RNA was of high quality (RIN∼10); the input mass was 100 ng. Libraries were prepared according to manufacturer’s instructions; the NEBNext rRNA Depletion Kit v2 was applied to remove human ribosomal RNA before strand-specific libraries were constructed using the NEBNext Ultra II Directional RNA Library Prep Kit for Illumina (E7760). The libraries were indexed with custom adapters and barcode tags (including dual indexing(Lamble et al. 2013)) and sequenced on an Illumina NovaSeq6000 v1.5 in 150bp paired-end mode. A minimum of 80 million reads were obtained per sample.

### Human transcriptome analysis

Raw Fastq files were trimmed for the presence of Illumina adapter sequences using Cutadapt v1.2.1 (Martin 2011). Reads were aligned to hg19 transcripts (*n*=180,253) using Kallisto v0.46.0 (Bray et al. 2016) to produce gene-level TPM estimates by aggregating transcripts per gene. Differential expression analysis was performed using edgeR (Robinson et al. 2010). Read counts were first TMM normalised (Robinson and Oshlack 2010) to account for differences in library size. Differentially expressed genes were identified by genewise exact test (Robinson and Smyth 2008); only genes which were significant after Benjamini-Hochberg multiple testing correction were reported as differentially expressed.

### Viral transcriptome analysis

The raw fastQ files were filtered for viral reads using ReadItAndKeep (Hunt et al. 2022) and the files of viral reads produced were then used for downstream analysis of Subgenomic RNA. Briefly, ReadItAndKeep removes host reads from viral sequencing data by aligning all reads against a target genome using minimap2 and retaining only reads that match at least 50bp or 50% of the length of the read. This approach has been shown to have 100% sensitivity and 99.894% specificity in distinguishing human from viral reads with illumina sequencing. ReadItAndKeep version 0.1.0 was downloaded from bioconda and run using the Wuhu-Hu-1 genome (NCBI Reference Sequence: NC_045512.2, download from www.ncbi.nlm.nih.gov/nuccore/NC_045512) as the target sequence the paired illumina reads in fastq format. The percentage of viral reads was calculated as the number of reads retained by ReadItAndKeep divided by the total number of reads.

To calculate the proportion of genomic:subgenomic reads we measured read-depth over SARS-CoV-2 Orf1 which is present in all genomic RNAs and none of the canonical subgenomic RNAs, to the read-depth over Orf10 which is present in all canonical subgenomic RNAs (Long 2021). A limitation of our method is that we do not consider low frequency non-canonical subgenomic RNAs.

## Figure Legends

**Supplementary Figure 1. CRISPR-editing with sgRNA targeted against *EXOSC2* in Calu-3 cells**. (**a**) Sanger sequencing traces demonstrating spCas9 cut site adjacent to PAM and subsequent waveform decomposition in *EXOSC2* edited cells. (**b**) Indel distribution of *EXOSC2* edited Calu-3 cells. (**c**) qPCR reveals that expression of *EXOSC2* mRNA is reduced in edited Calu-3 cells; comparison is made with control CRISPR editing targeting *HPRT*. (**d**) Immunoblotting for EXOSC2 protein revealed reduced expression in EXOSC2 edited Calu-3 cells compared to HPRT edited cells and WT unedited cells; recovery of EXOSC2 expression was achieved with EXOSC2 reconstitution. GAPDH served as a loading control and a serial dilution of protein from WT cells is shown for reference.

**Supplementary Figure 2: Transcriptomic analysis of the effect of SARS-CoV-2 infection and/or reduced EXOSC2 expression in Calu-3 cells**. Volcano plot to compare gene expression changes coincident with SARS-CoV-2 infection in (**a**) WT unedited Calu-3 cells and (**b**) in cells with prior CRISPR editing of EXOSC2. (**c**) Volcano plot to compare gene expression in SARS-CoV-2 infected Calu-3 cells in the presence and absence of CRISPR editing of EXOSC2. Dotted lines represent fold change of +/- 2 and a Bonferroni multiple testing threshold for p-value by genewise exact test.

**Supplementary Table 1. Association of lung expression with risk for clinical COVID-19 for genes encoding 332 host proteins which interact with SARS-CoV-2 proteins**. For each gene a set of SNPs was identified which were association with expression of the gene in lung tissue (GTEx v7) (Lonsdale et al. 2013). For each of these eQTLs we measured association with risk of clinical COVID-19 (COVID-19 Host Genetics Initiative 2021). The table details the lead SNP by association with lung expression and with COVID-19 risk. We tested for an association of gene expression with COVID-19 risk using FUSION (Gusev et al. 2016); the total number of SNPs in each test and the test statistics are detailed.

## References

Allmang C, Mitchell P, Petfalski E, Tollervey D. 2000. Degradation of ribosomal RNA precursors by the exosome. Nucleic Acids Res 28: 1684–1691.

Bray NL, Pimentel H, Melsted P, Pachter L. 2016. Near-optimal probabilistic RNA-seq quantification. Nat Biotechnol 34: 525–527.

Brodin P. 2021. Immune determinants of COVID-19 disease presentation and severity. Nat Med 27: 28–33.

Cagno V. 2020. SARS-CoV-2 cellular tropism. Lancet Microbe 1: e2–e3.

Choi UY, Kang J-S, Hwang YS, Kim Y-J. 2015. Oligoadenylate synthase-like (OASL) proteins: dual functions and associations with diseases. Exp Mol Med 47: e144.

Chu H, Chan JF-W, Yuen TT-T, Shuai H, Yuan S, Wang Y, Hu B, Yip CC-Y, Tsang JO-L, Huang X, et al. 2020. Comparative tropism, replication kinetics, and cell damage profiling of SARS-CoV-2 and SARS-CoV with implications for clinical manifestations, transmissibility, and laboratory studies of COVID-19: an observational study. Lancet Microbe 1: e14–e23.

Conant D, Hsiau T, Rossi N, Oki J, Maures T, Waite K, Yang J, Joshi S, Kelso R, Holden K, et al. 2022. Inference of CRISPR Edits from Sanger Trace Data. CRISPR J 5: 123–130.

COVID-19 Host Genetics Initiative. 2021. Mapping the human genetic architecture of COVID-19. Nature. http://dx.doi.org/10.1038/s41586-021-03767-x.

Darby AC, Hiscox JA. 2021. Covid-19: variants and vaccination. BMJ 372: n771.

Dong E, Du H, Gardner L. 2020. An interactive web-based dashboard to track COVID-19 in real time. Lancet Infect Dis 20: 533–534.

Eckard SC, Rice GI, Fabre A, Badens C, Gray EE, Hartley JL, Crow YJ, Stetson DB. 2014. The SKIV2L RNA exosome limits activation of the RIG-I-like receptors. Nat Immunol 15: 839–845.

Fathi M, Vakili K, Sayehmiri F, Mohamadkhani A, Hajiesmaeili M, Rezaei-Tavirani M, Eilami O. 2021. The prognostic value of comorbidity for the severity of COVID-19: A systematic review and meta-analysis study. PLoS One 16: e0246190.

Gordon DE, Jang GM, Bouhaddou M, Xu J, Obernier K, White KM, O’Meara MJ, Rezelj VV, Guo JZ, Swaney DL, et al. 2020. A SARS-CoV-2 protein interaction map reveals targets for drug repurposing. Nature 583: 459–468.

Gunst JD, Staerke NB, Pahus MH, Kristensen LH, Bodilsen J, Lohse N, Dalgaard LS, Brønnum D, Fröbert O, Hønge B, et al. 2021. Efficacy of the TMPRSS2 inhibitor camostat mesilate in patients hospitalized with Covid-19-a double-blind randomized controlled trial. EClinicalMedicine 35: 100849.

Gusev A, Ko A, Shi H, Bhatia G, Chung W, Penninx BWJH, Jansen R, de Geus EJC, Boomsma DI, Wright FA, et al. 2016. Integrative approaches for large-scale transcriptome-wide association studies. Nat Genet 48: 245–252.

Hillen HS, Kokic G, Farnung L, Dienemann C, Tegunov D, Cramer P. 2020. Structure of replicating SARS-CoV-2 polymerase. Nature 584: 154–156.

Hoffmann M, Kleine-Weber H, Schroeder S, Krüger N, Herrler T, Erichsen S, Schiergens TS, Herrler G, Wu N-H, Nitsche A, et al. 2020. SARS-CoV-2 Cell Entry Depends on ACE2 and TMPRSS2 and Is Blocked by a Clinically Proven Protease Inhibitor. Cell 181: 271–280.e8.

Ho JSY, Zhu Z, Marazzi I. 2021. Unconventional viral gene expression mechanisms as therapeutic targets. Nature 593: 362–371.

Holshue ML, DeBolt C, Lindquist S, Lofy KH, Wiesman J, Bruce H, Spitters C, Ericson K, Wilkerson S, Tural A, et al. 2020. First Case of 2019 Novel Coronavirus in the United States. N Engl J Med 382: 929–936.

Huffman JE, Butler-Laporte G, Khan A, Pairo-Castineira E, Drivas TG, Peloso GM, Nakanishi T, COVID-19 Host Genetics Initiative, Ganna A, Verma A, et al. 2022. Multi-ancestry fine mapping implicates OAS1 splicing in risk of severe COVID-19. Nat Genet. http://dx.doi.org/10.1038/s41588-021-00996-8.

Hunt M, Swann J, Constantinides B, Fowler PW, Iqbal Z. ReadItAndKeep: rapid decontamination of SARS-CoV-2 sequencing reads. http://dx.doi.org/10.1101/2022.01.21.477194.

Iida K, Ajiro M, Muramoto Y, Takenaga T, Denawa M, Kurosawa R, Noda T, Hagiwara M. 2021. Switching of OAS1 splicing isoforms mitigates SARS-CoV-2 infection. bioRxiv 2021.08.23.457314. https://www.biorxiv.org/content/10.1101/2021.08.23.457314 (Accessed March 4, 2022).

Jayk Bernal A, Gomes da Silva MM, Musungaie DB, Kovalchuk E, Gonzalez A, Delos Reyes V, Martín-Quirós A, Caraco Y, Williams-Diaz A, Brown ML, et al. 2021. Molnupiravir for Oral Treatment of Covid-19 in Nonhospitalized Patients. N Engl J Med. http://dx.doi.org/10.1056/NEJMoa2116044.

Kilchert C, Wittmann S, Vasiljeva L. 2016. The regulation and functions of the nuclear RNA exosome complex. Nat Rev Mol Cell Biol 17: 227–239.

Lamble S, Batty E, Attar M, Buck D, Bowden R, Lunter G, Crook D, El-Fahmawi B, Piazza P. 2013. Improved workflows for high throughput library preparation using the transposome-based Nextera system. BMC Biotechnol 13: 104.

Long S. 2021. SARS-CoV-2 Subgenomic RNAs: Characterization, Utility, and Perspectives. Viruses 13. http://dx.doi.org/10.3390/v13101923.

Lonsdale J, Thomas J, Salvatore M, Phillips R, Lo E, Shad S, Hasz R, Walters G, Garcia F, Young N, et al. 2013. The Genotype-Tissue Expression (GTEx) project. Nat Genet 45: 580–585.

Manjili RH, Zarei M, Habibi M, Manjili MH. 2020. COVID-19 as an Acute Inflammatory Disease. J Immunol 205: 12–19.

Martin M. 2011. Cutadapt removes adapter sequences from high-throughput sequencing reads. EMBnet.journal 17: 10–12.

Menni C, Valdes AM, Freidin MB, Sudre CH, Nguyen LH, Drew DA, Ganesh S, Varsavsky T, Cardoso MJ, El-Sayed Moustafa JS, et al. 2020. Real-time tracking of self-reported symptoms to predict potential COVID-19. Nat Med 26: 1037–1040.

Molleston JM, Cherry S. 2017. Attacked from All Sides: RNA Decay in Antiviral Defense. Viruses 9. http://dx.doi.org/10.3390/v9010002.

Mullani N, Porozhan Y, Costallat M, Batsché E, Goodhardt M, Cenci G, Mann C, Muchardt C. 2021 Reduced RNA turnover as a driver of cellular senescence. Life Science Alliance doi: 10.26508/lsa.202000809.

Olliaro P, Torreele E, Vaillant M. 2021. COVID-19 vaccine efficacy and effectiveness—the elephant (not) in the room. The Lancet Microbe 2: e279–e280.

Pouwels KB, Pritchard E, Matthews P, Stoesser NB, Eyre DW, Vihta K-D, House T, Hay J, Bell J, Newton J, et al. 2021. Impact of Delta on viral burden and vaccine effectiveness against new SARS-CoV-2 infections in the UK. bioRxiv. http://medrxiv.org/lookup/doi/10.1101/2021.08.18.21262237.

Rialdi A, Hultquist J, Jimenez-Morales D, Peralta Z, Campisi L, Fenouil R, Moshkina N, Wang ZZ, Laffleur B, Kaake RM, et al. 2017. The RNA Exosome Syncs IAV-RNAPII Transcription to Promote Viral Ribogenesis and Infectivity. Cell 169: 679–692.e14.

Robinson MD, McCarthy DJ, Smyth GK. 2010. edgeR: a Bioconductor package for differential expression analysis of digital gene expression data. Bioinformatics 26: 139–140.

Robinson MD, Oshlack A. 2010. A scaling normalization method for differential expression analysis of RNA-seq data. Genome Biol 11: R25.

Robinson MD, Smyth GK. 2008. Small-sample estimation of negative binomial dispersion, with applications to SAGE data. Biostatistics 9: 321–332.

Schoggins JW, Wilson SJ, Panis M, Murphy MY, Jones CT, Bieniasz P, Rice CM. 2011. A diverse range of gene products are effectors of the type I interferon antiviral response. Nature 472: 481–485.

Shilo S, Rossman H, Segal E. 2021. Signals of hope: gauging the impact of a rapid national vaccination campaign. Nat Rev Immunol 21: 198–199.

The RECOVERY Collaborative Group. 2021. Dexamethasone in Hospitalized Patients with Covid-19. N Engl J Med 384: 693–704.

Tomecki R, Kristiansen MS, Lykke-Andersen S, Chlebowski A, Larsen KM, Szczesny RJ, Drazkowska K, Pastula A, Andersen JS, Stepien PP, et al. 2010. The human core exosome interacts with differentially localized processive RNases: hDIS3 and hDIS3L. EMBO J 29: 2342–2357.

Tyanova S, Temu T, Cox J. 2016. The MaxQuant computational platform for mass spectrometry-based shotgun proteomics. Nat Protoc 11: 2301–2319.

Viswanathan T, Arya S, Chan S-H, Qi S, Dai N, Misra A, Park J-G, Oladunni F, Kovalskyy D, Hromas RA, et al. 2020. Structural basis of RNA cap modification by SARS-CoV-2. Nat Commun 11: 3718.

Weston S, Baracco L, Keller C, Matthews K, McGrath ME, Logue J, Liang J, Dyall J, Holbrook MR, Hensley LE, et al. 2020. The SKI complex is a broad-spectrum, host-directed antiviral drug target for coronaviruses, influenza, and filoviruses. Proc Natl Acad Sci U S A 117: 30687–30698.

Wickenhagen A, Sugrue E, Lytras S, Kuchi S, Noerenberg M, Turnbull ML, Loney C, Herder V, Allan J, Jarmson I, et al. 2021. A prenylated dsRNA sensor protects against severe COVID-19. Science 374: eabj3624.

Wyler E, Mösbauer K, Franke V, Diag A, Gottula LT, Arsiè R, Klironomos F, Koppstein D, Hönzke K, Ayoub S, et al. 2021. Transcriptomic profiling of SARS-CoV-2 infected human cell lines identifies HSP90 as target for COVID-19 therapy. iScience 24: 102151.

Zhang S, Cooper-Knock J, Weimer AK, Harvey C, Julian TH, Wang C, Li J, Furini S, Frullanti E, Fava F, et al. 2021. Common and rare variant analyses combined with single-cell multiomics reveal cell-type-specific molecular mechanisms of COVID-19 severity. medRxiv. http://dx.doi.org/10.1101/2021.06.15.21258703.

